# Altered hippocampal interneuron activity precedes ictal onset

**DOI:** 10.1101/064956

**Authors:** Mitra L. Miri, Martin Vinck, Rima Pant, Jessica A. Cardin

**Affiliations:** Department of Neuroscience, Yale University School of Medicine, New Haven CT Kavli Institute, Yale University, New Haven CT

**Keywords:** Seizure, interneuron, parvalbumin, somatostatin, optogenetics, channelrhodopsin, inhibition, hippocampus, local field potential, ictal

## Abstract

Although failure of GABAergic inhibition is a commonly hypothesized mechanism underlying seizure disorders, the series of events that precipitate a rapid shift from healthy to ictal activity remain unclear. Furthermore, the diversity of inhibitory interneuron populations poses a challenge for understanding local circuit interactions during seizure initiation. Using a combined optogenetic and electrophysiological approach, we examined the activity of two identified hippocampal interneuron classes during seizure induction in vivo. We identified cell type-specific differences in preictal firing patterns and input sensitivity of parvalbumin- and somatostatin-expressing interneurons. Surprisingly, the impact of both sources of inhibition remained intact throughout the preictal period and into the early ictal phase. Our findings suggest that the onset of ictal activity is not due to a failure of inhibition, but is instead associated with a decoupling of inhibitory cells from their normal relationship with the local hippocampal network.

## Introduction

Seizure activity is commonly considered to arise from an imbalance of excitation and inhibition in vulnerable neural circuits, leading to unconstrained activity that self-organizes into 57 patterns of hypersynchrony. One mechanism of such an imbalance may be a failure of GABAergic inhibition (Ziburkus et al., 2006). Acute blockade of GABA receptors rapidly initiates seizure activity (Rose and Blakemore, 1974; Treiman, 2001), suggesting the necessity of synaptic inhibition to maintain healthy activity patterns. Previous work has also highlighted the potential preictal contribution of excitatory GABAergic effects due to Cl-accumulation (Cossart et al., 2005; Palma et al., 2006; Miles et al., 2012) or loss of inhibition due to depletion of GABAergic release (Zhang et al., 2012). Developmental dysregulation of inhibitory interneurons causes chronic epileptic disorders (Lau et al., 2000; Rossignol et al., 2013; Tai et al., 2014), and interneurons also appear to be particularly sensitive to seizure-related damage (Sloviter, 1987; de Lanerolle et al., 1989; Robbins et al., 1991; Rice et al., 1996; Gibbs et al., 1997; Cossart et al., 2001). However, the circuit mechanisms underlying seizure initiation and the specific role of GABAergic interneurons remain largely unknown.

Work from *in vitro* and *in vivo* animal models has suggested that different neural populations may have distinct preictal roles in seizure initiation. Excitatory and inhibitory neuron activity has been found to increase preictally (Timofeev et al., 2002; Bower and Buckmaster, 2008; Cymerblit-Sabba and Schiller, 2010; Jiruska et al., 2010; Grasse et al., 2013; Toyoda et al., 2015) or to be modulated in opposition before ictal onset (Ziburkus et al., 2006). Furthermore, recent work using single-unit recordings in human patients found stronger preictal activation of putative interneuron populations than other cells (Truccolo et al., 2011). However, one challenge in examining the respective roles of excitatory and inhibitory cells in seizure initiation is the diversity of inhibitory interneurons. Neocortical GABAergic interneurons exhibit a wide variety of morphologies, molecular markers, and activity patterns, and make synapses on different subcellular domains of target pyramidal cells (Rudy et al., 2011). In the hippocampus, these include the soma-targeting, axo-axonic and basket cells that co-express the calcium binding protein parvalbumin (PV) and the dendrite-targeting O-LM and bistratified interneurons that co-express the peptide somatostatin (SOM) (Buhl et al., 1994; Sik et al., 1995; Klausberger et al., 2003; Petilla Interneuron Nomenclature et al., 2008; Lapray et al., 2012). Despite their diverging cellular properties, these cell classes are difficult to identify *in vivo* using traditional recording methods.

Here we used optogenetic tools to identify, track, and probe two distinct populations of hippocampal interneurons, the PV- and SOM-expressing cells, in two models of acute seizure initiation *in vivo*. We find cell type-specific differences in the preictal activity of PV and SOM cells and in the evolution of their sensitivity to input. However, the inhibitory influence of interneuron firing on nearby neurons remains largely intact throughout the preictal and early ictal periods, suggesting that seizure does not arise from a failure of GABAergic inhibition.

## Results

To examine the preictal activity of identified hippocampal interneurons, we performed tetrode recordings of isolated single units and local field potentials (LFPs) from hippocampal CA1 in anesthetized mice expressing Channelrhodopsin-2 (ChR2) in target cells. Seizure activity was induced with systemic administration of the chemoconvulsant Pentylenetetrazol (PTZ) (Figure 1A). During the baseline period, we identified PV- (n = 56 cells in 45 mice) and SOM- (n = 42 cells in 34 mice) expressing interneurons in PV-Cre/ChR2 or SOM-Cre/ChR2 mice, respectively, by their short-latency, low-jitter responses to blue light (Figure 1B; see Methods) (Cardin et al., 2009; Lima et al., 2009). On average, PV cells displayed narrow spike waveforms (Figures 1C and S1), whereas SOM cells exhibited a broader waveform distribution (Figure S1A). However, there was extensive overlap of waveform characteristics among the populations (Figure S1), indicating that spike waveform alone is not sufficient to distinguish cell types under these conditions. Histological analysis confirmed that the two identified interneuron populations were largely non-overlapping within CA1 (Figure S2). In a subset of experiments, unidentified cells (n = 49 cells in 26 mice) were simultaneously recorded along with ChR2-tagged units in PV-Cre and SOM-Cre mice or in the pyramidal cell layer of wild-type mice (Figure S1). A subset of these unidentified cells (n = 26 cells in 16 mice) were regular spiking (RS), putative excitatory cells with characteristic broad spike waveforms and relatively low baseline firing rates (Figures 1C and S1).

**Fig. 1.**
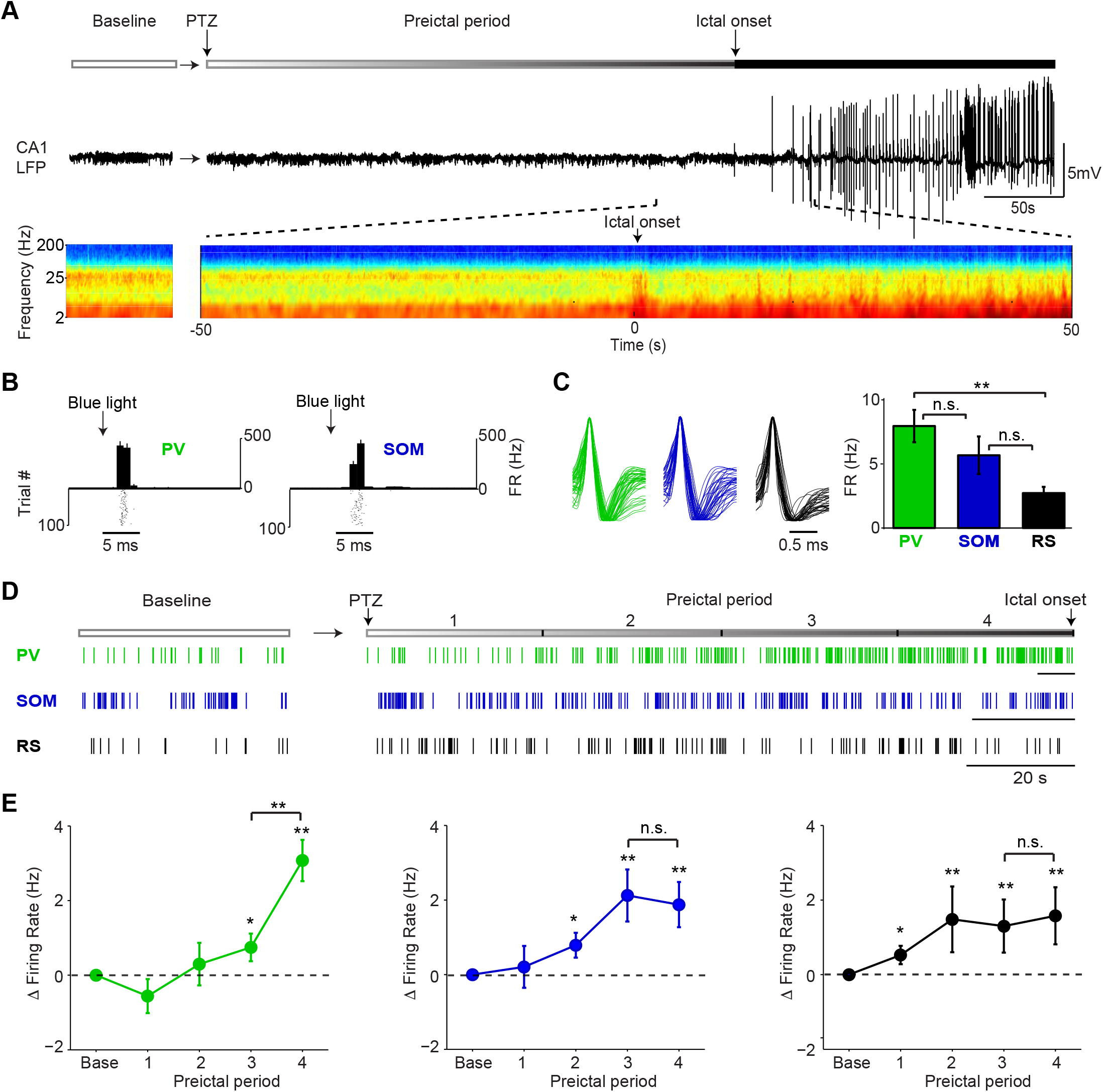
Firing rate changes of ChR2-tagged interneurons precede seizure initiation. **(A)** Upper: Schematic of experimental paradigm showing baseline, PTZ injection and preictal 265 period leading up to ictal onset. Middle: LFP trace from hippocampal CA1 recording. Lower: Spectrogram showing LFP power as a function of time from ictal onset (x-axis) and frequency (y-axis; shown on log-10 scale). **(B)** Peri-pulse time-histogram around 5ms laser pulses together with raster plot during baseline for example PV- and SOM-expressing interneurons in PV-Cre/ChR2 and SOM-Cre/ChR2 mice, respectively. **(C)** Left: Averaged, normalized action potential waveforms for all recorded PV, SOM and putative RS cells. Right: Mean baseline firing rates (Hz) for PV, SOM and RS cells recorded from hippocampal CA1. **(D)** Spike trains for example PV, SOM and RS cells. All horizontal scale bars correspond to 20s. Note that cells were recorded from different animals that each had different latencies to ictal onset. **(E)** Mean changes in firing rate as compared to baseline over four preictal periods for PV (n = 18 cells in 13 mice), SOM (n = 16 cells in 10 mice) and RS (n = 20 cells in 12 mice) populations. Note that pairwise statistical comparisons are only shown between 3^rd^ and 4^th^ preictal period. Error bars denote mean s.e.m. ∗p<0.05, ∗∗p<0.01.

In an initial series of experiments, we assessed the spontaneous activity of these three cell classes during a baseline period and four preictal periods of equal duration leading into PTZ-induced seizure (Figure 1D). PV, SOM and RS cells exhibited increased firing rates following PTZ administration, but showed markedly different firing rate trajectories (Figure 1E). Strikingly, we found that most PV cells strongly increased their firing rate in the last preictal period as compared to the first (94.4%, p<0.001, Binomial test) or third (83.3%, p<0.001; Figure S3). In contrast, this progressive, late increase in firing rate was not observed in SOM or RS cells (Figures 1E and S3). Increased PV cell firing was independent of the latency to ictal onset (Figure S4) and was observed in the absence of significant changes in spike waveform amplitude over time (Figure S5). We next explored whether preictal firing rate changes were 122 accompanied by changes in the temporal spike pattern. Immediately preceding ictal onset, PV, but not SOM or RS, cell firing became significantly more regular (i.e., less bursty) (Figure 2A). In addition, PV cells showed an increased tendency to fire spikes separated by short (<10ms) 125 inter-spike-intervals (ISI; Figure S6). Unidentified cells with narrow spikes did not exhibit changes in firing rate, firing regularity, or ISI statistics (Figure S7).

**Fig 2.**
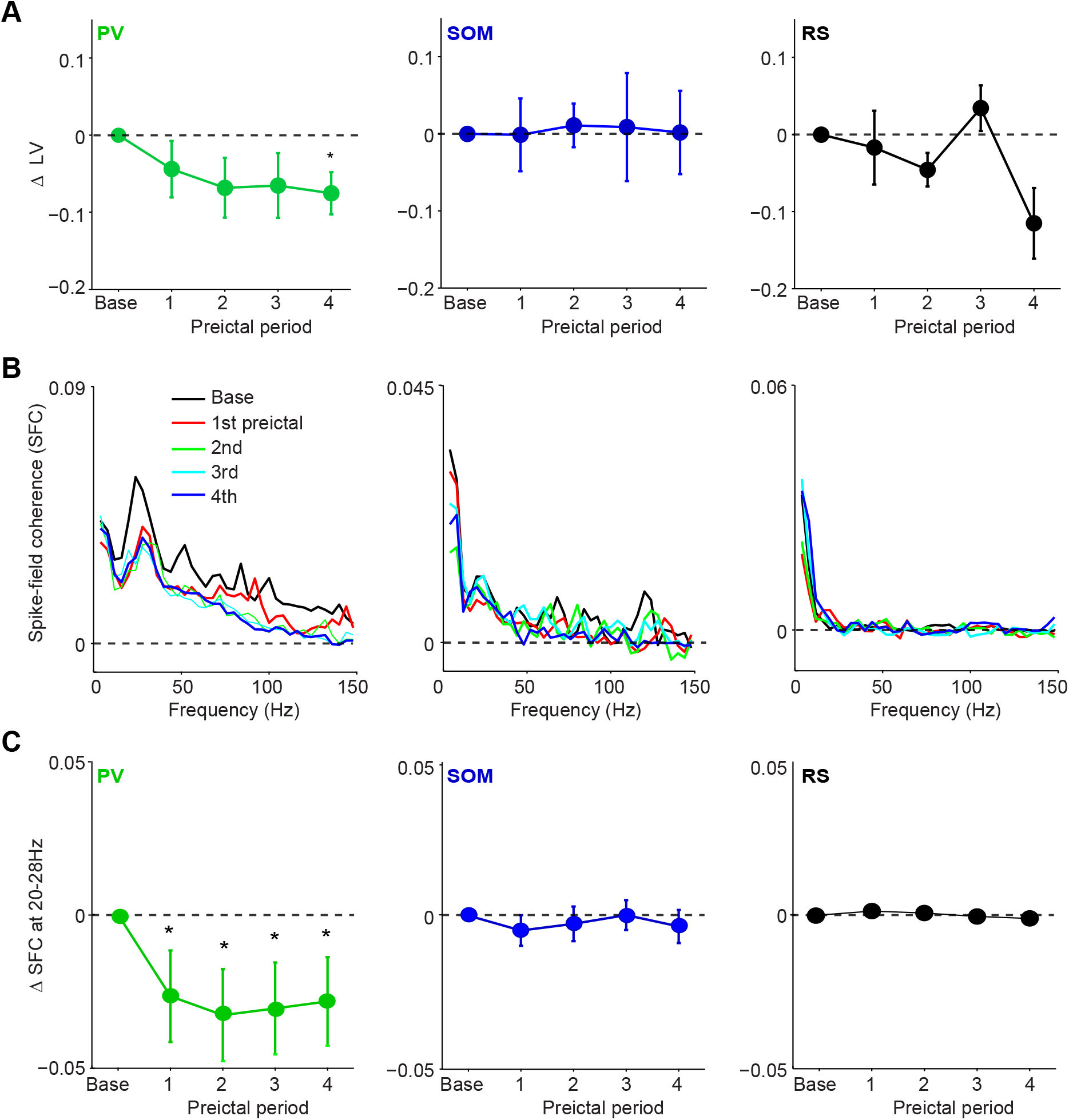
Preictal changes in temporal patterning of interneuron activity. **(A)** Mean changes in local variation (LV) of firing as compared to baseline over four preictal periods for PV, SOM and RS cells during acute PTZ seizure. LV is a measure of firing irregularity, where LV>1 indicates irregular firing (see Methods). **(B)** Left: Mean spike-field coherence (SFC) as a function of frequency (Hz) during baseline and four preictal periods for PV-expressing cells. Middle and right: Same for SOM and RS cells, respectively. **(C)** Mean changes in SFC in 20-28 Hz band as compared to baseline over four preictal periods for PV, SOM and RS cells. Error bars denote mean S s.e.m. ∗p<0.05.

In a separate series of experiments, we observed a similar increase in PV cell firing rates during the late preictal period preceding Pilocarpine-induced seizures (Figure S8), suggesting that this is not a unique feature of PTZ-induced seizures but rather may be a general feature of preictal activity in CA1. PV cells also showed increased regularity and shorter ISIs during preictal periods preceding Pilocarpine-induced seizures, in the absence of changes in spike waveform amplitude (Figure S8D).

The progressive changes in PV interneuron firing rate and temporal pattern suggest that the relationship between these cells and the surrounding local network may be altered prior to seizure initiation. We therefore computed the mean spike field coherence (SFC) for PV, SOM and RS cells during baseline activity and across the four preictal periods (Figure 2B). We observed a prominent peak in the 20-28Hz band of the LFP (Chen et al., 2011; Cabral et al., 2014; Sauer et al., 2015) under baseline and preictal conditions (Fig. 1A). PV cells showed a peak in SFC in the 20-28Hz frequency band that decreased significantly following PTZ administration (Figure 2B-C) in the absence of any loss in LFP power or changes in SFC in the theta or high gamma bands, other prominent hippocampal rhythms associated with interneuron activity (Figure S9). In contrast, neither SOM nor RS cells showed a change in SFC across the preictal periods, suggesting a specific decoupling of PV cells from their normal temporal relationship with the local hippocampal network during the onset of ictal activity.

Together, these findings highlight cell type-specific changes in PV interneuron activity leading up to the onset of ictal activity. To examine whether preictal changes in interneuron output were accompanied by changes in sensitivity to input, we tested the responses of PV and SOM cells to optogenetic stimulation during each preictal period. We measured the probability of interneuron spiking in response to light pulses of varying intensity (Figure 3A). PV cells showed no progressive change in the slope or maximal response (Rmax) of the input-output function (Figure 3B-C). In contrast, SOM cells showed a significant increase in slope and a significant decrease in Rmax across the preictal periods, suggesting a progressive preictal alteration in their sensitivity to inputs.

**Fig. 3.**
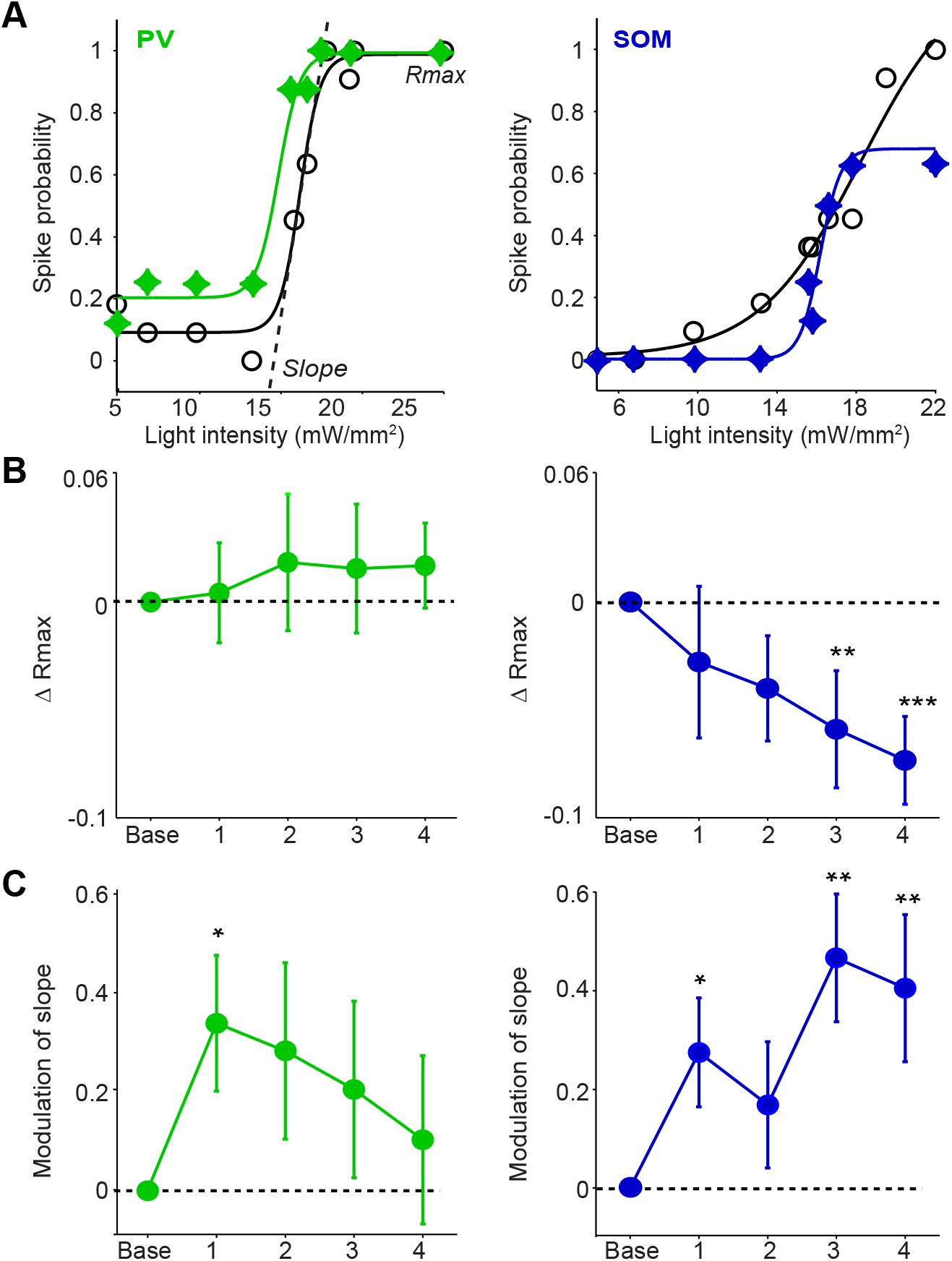
Preictal changes in interneuron sensitivity to inputs. **(A)** Response curve of spike probability as a function of light pulse intensity (mW/mm^2^) with sigmoid curve fit during baseline (black) and 4^th^ preictal period (colors) for example PV (left) and SOM (right) cells. **(B)** Mean change in Rmax compared to baseline (dotted line) over four preictal periods for PV (n =17 cells in 15 mice) and SOM (n=21 cells in 20 mice) cells. **(C)** Mean modulation of the slope parameter compared to baseline over four preictal periods for PV and SOM cells. **(B-C)** Significance to baseline: ∗p<0.05, ∗∗p<0.01, ∗∗∗p<0.001. Error bars denote mean s s.e.m.

To assay whether the observed changes in interneuron activity were associated with altered inhibition of their targets, we measured the impact of ChR2-evoked interneuron spiking on the firing rate of nearby RS cells. During baseline activity, we found that the firing rate of RS cells decreased following ChR2-evoked PV and SOM cell spiking (Figure 4A-B). We compared the impact of ChR2-evoked inhibition during the four preictal periods and an additional period immediately following ictal onset. RS firing suppression was not significantly changed across the preictal and early ictal periods as compared to baseline when the PV cells were driven at either moderate or high light intensities (see Methods; Figure 4C-D). RS suppression by SOM cell spiking was likewise maintained throughout preictal and early ictal periods (Figure 4E-F). Thus, both PV- and SOM-mediated inhibition appear largely intact preceding ictal onset and remain so during the transition to ictal activity.

**Fig 4.**
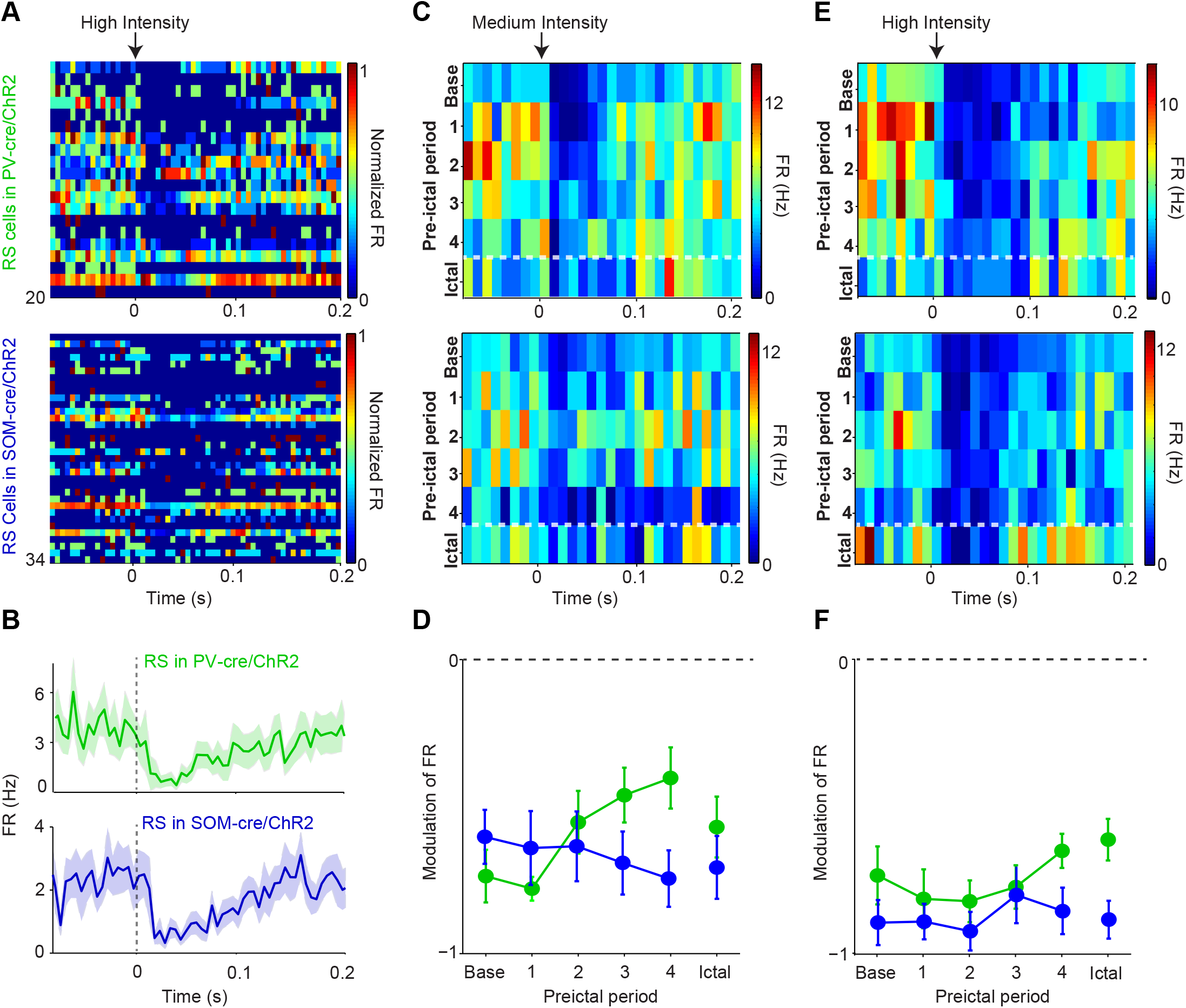
Intact preictal suppression of RS firing by evoked inhibition. **(A)** Normalized firing rate (FR) as a function of time (s) from laser pulse onset (t=0) for RS cells recorded simultaneously with PV cells in PV-Cre/ChR2 mice (n = 20 cells in 12 mice) or recorded with SOM cells in SOM-Cre/ChR2 mice (n = 34 cells in 16 mice) during baseline activity. For purposes of visualization, firing rates were normalized to the maximum firing rate for each cell. **(B)** Mean changes in baseline firing rate (Hz) of RS cells as a function of time (s) from laser pulse onset (dashed grey line). **(C)** Upper: Average firing rate of RS cells (n =11 cells in 8 mice) recorded during PV/ChR2 experiments as a function of time (s) from laser pulse onset during baseline, the four preictal periods and a 60s period following ictal onset after PTZ administration (see Methods). Lower: Average firing rate of RS cells (n =11 cells in 7 mice) recorded during SOM/ChR2 experiments as a function of time (s) from laser pulse onset during baseline, the four preictal periods and following ictal onset. For these plots, we considered only light pulses of medium light intensity (see Methods). **(D).** Mean modulation of firing rate (see Methods) after medium intensity laser pulse (0 to 50ms) compared to pre-pulse (-200 to 0ms) firing rate for RS cells recorded during PV/ChR2 (green) and SOM/ChR2 (blue) experiments.Modulation scores for RS in PV-Cre and RS in SOM-Cre were significantly different from zero for all periods. **(E).** As in C, but for high intensity light pulses. **(F)** As in D, but for high intensity light pulses (see Methods). No significant changes were observed in modulation scores between periods for medium or high intensity pulses for either group. Error bars denote mean d s.e.m.

At the ictal transition, changes in interneuron activity mainly appeared to be at the level of firing rates, rather than postsynaptic impact on nearby RS cells. We next examined whether interneuron firing remained elevated during the ictal period. Rigorous spike waveform identification after ictal onset is highly challenging. However, we were able to track a subset of recorded neurons through an initial 60-second ictal period. We found that after ictal onset, PV and SOM firing decreased from the preictal peak back to baseline levels (Figure 5A, B). In contrast, RS cells continued to fire at elevated rates (Figure 5C), suggesting a sustained decoupling of excitatory and inhibitory spiking.

**Fig 5.**
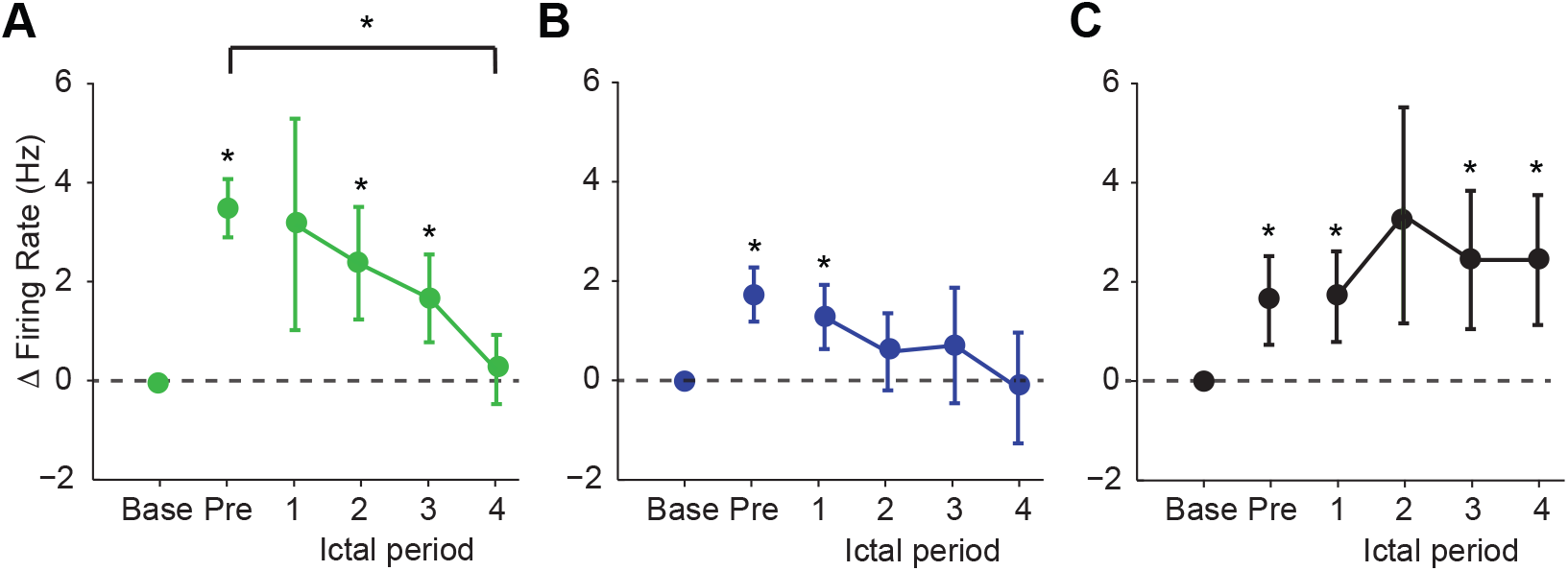
Interneuron firing rates decrease after ictal onset. **(A)** Mean changes in firing rate as compared to baseline during four ictal periods for PV interneurons (n = 9 cells in 8 mice). Firing rate in fourth ictal period was significantly lower than in fourth pre-ictal period for PV cells (p<0.05). **(B)** Mean ictal changes in SOM interneuron firing rate (n = 18 cells in 17 mice). **(C)** Mean ictal changes in RS cell firing rate (n = 16 cells in 9 mice). Error bars denote mean s.e.m. ∗p<0.05, ∗∗p<0.01

## Discussion

During spontaneous and evoked neural activity, excitation and inhibition are tightly coupled in amplitude (Shu et al., 2003; Haider et al., 2006; Xue et al., 2014) and temporal pattern (Pouille and Scanziani, 2001; Wehr and Zador, 2003; Higley and Contreras, 2006; Cardin et al., 2010). Disruption of the excitatory-inhibitory balance profoundly alters circuit function (Fritschy, 2008; Ziburkus et al., 2013). We found that PV- and SOM-expressing interneurons and RS, putative excitatory cells exhibited different preictal firing rate trajectories.SOM and RS cells showed an early, modest increase in firing, whereas PV cells exhibited a late increase in firing immediately before ictal onset. During the early ictal period, PV and SOM cell firing rates returned to baseline levels, whereas RS cell firing remained elevated. These changes in PV and SOM cell firing rates occurred in the absence of changes in the impact of ChR2-evoked inhibition on local RS cells. Overall, these results suggest that seizure did not arise from either an overall failure of GABAergic inhibition or from unconstrained excitation, but rather from a decoupling of excitatory and inhibitory activity.

Fast-spiking, PV-expressing interneurons are coupled to both high-frequency and theta rhythms in the hippocampus, and are thought to contribute to the fine temporal organization of 189 excitatory-inhibitory interactions (Buzsaki et al., 2003; Csicsvari et al., 2003; Klausberger et al., 2003; Forro et al., 2015). The earliest preictal change we observed was a loss of the strong spike-field coherence of PV cell spiking to a prominent low-gamma band previously observed in mouse hippocampus (Chen et al., 2011; Cabral et al., 2014; Sauer et al., 2015). Thus, despite their increase in firing rate and decreased burstiness at the transition to ictal activity, PV interneurons were already decoupled from their precise temporal relationship with the surrounding local network prior to seizure initiation. Increased PV interneuron activity could lead to irregular firing patterns or bursts, potentially resulting in hyper-synchronized, pathological entrainment of excitatory activity (Yekhlef et al., 2015). Previous work has suggested that increased inhibitory synaptic activity or the onset of depolarization block in fast-spiking interneurons could precipitate seizure onset (Velazquez and Carlen, 1999; Timofeev et al., 2002; Fujiwara-Tsukamoto et al., 2007; Gnatkovsky et al., 2008). We found that the onset of ictal spikes coincided with a sharp increase in PV interneuron spiking. However, we found no evidence for decreased interneuron spike amplitude, suggesting that these interneurons did not enter depolarization block prior to ictal onset. In contrast to the PV interneurons, SOM 204 interneurons showed an earlier increase in firing rate but no change in the temporal pattern of their spiking. SOM, but not PV, cells exhibited altered sensitivity to input and a decreased range of output responses during the late preictal period. This diminished dynamic range could compromise the ability of SOM cells to participate in balanced excitatory-inhibitory interactions within the local hippocampal circuit (Lovett-Barron et al., 2012; Royer et al., 2012).

Although PV and SOM interneuron populations showed increased activity during the preictal period, these changes were transient and not accompanied by reduced RS activity.Early preictal decoupling of PV cells from normal hippocampal rhythms was followed by a transient late preictal rise in PV interneuron firing that was not accompanied by a change in RS cell firing. In the early ictal period PV and SOM firing rates decreased, whereas RS activity remained stable at an elevated firing rate. The activity of interneuron and excitatory neuron populations was thus progressively decoupled in multiple ways during the transition to seizure.

Previous work has found extensive heterogeneity in preictal firing rate trajectories of extracellularly recorded putative excitatory and inhibitory neurons. Intracellular and extracellular recordings of fast-spiking, putative PV interneurons in the cortex and hippocampus have found increased preictal firing rates (Timofeev et al., 2002; Gnatkovsky et al., 2008) or a transient increase in firing immediately preceding ictal onset (Grasse et al., 2013). Some reports suggest increased firing of regular spiking putative excitatory neurons in the entorhinal cortex and hippocampus (Cymerblit-Sabba and Schiller, 2010; Jiruska et al., 2010; Fujita et al., 2014), but others found a mix of increases and decreases within multiple hippocampal areas (Bower and Buckmaster, 2008; Toyoda et al., 2015). To our knowledge, there are no previous data on the preictal firing rate trajectories of identified SOM interneurons. We found that identification by action potential waveform or baseline firing rate was not a reliable indicator of cell identity for hippocampal PV interneurons. In addition, in agreement with previous work (Klausberger et al., 2003; Halabisky et al., 2006; Katona et al., 2014), we found that SOM interneurons exhibited varied action potential durations and could not be distinguished from regular spiking excitatory neurons by waveform. Optical tagging with Cre-dependent ChR2 allowed identification of each population despite overlapping action potential characteristics. The three cell classes we examined demonstrated distinct trajectories, suggesting that some of the preictal heterogeneity observed in other studies may arise from a mixed population of unidentified cells.

We examined interneuron activity in two models of acute induction of status epilepticus, which may provide insight into the mechanisms by which normal, healthy neural circuits transition to pathological patterns of activation. We used both PTZ, thought to be a competitive antagonist of the GABA_A_ receptor (Huang et al., 2001), and Pilocarpine, a nonselective muscarinic acetylcholine receptor agonist (Turski et al., 1989). Despite distinct pharmacological mechanisms, we found similar trajectories for PV interneuron activity in both models, suggesting that preictal increases in PV activity may be a common element of acute seizure initiation. Previous work on acute seizure induction likewise observed similar overall preictal firing rate trajectories across several models (Cymerblit-Sabba and Schiller, 2010). Because these drugs arrive rapidly in the brain at effective concentrations following systemic administration (Yonekawa et al., 1980; Ramzan and Levy, 1985), it is unlikely that the progression of interneuron firing changes from preictal to ictal periods resulted from gradual accumulation of chemoconvulsants in neural tissue. Our experiments were conducted under anesthesia to reduce movement artifacts and allow continued recording of small GABAergic neurons, and the presence of anesthestics could potentially prolong the preictal period and reduce seizure activity(Murao et al., 2002; Fang and Wang, 2015; Grover et al., 2016). However, the light anesthesia we used allowed normal spontaneous hippocampal firing patterns to be maintained and promoted short ictal onset delays.

Our data suggest that cell type-specific disruption of finely tuned interneuron relationships with the local hippocampal circuit may contribute to temporal reorganization of activity and decoupling of excitation and inhibition prior to seizure initiation. PV and SOM interneurons exhibited distinct profiles of preictal changes in firing and sensitivity to inputs. In particular, preictal disruption of PV interneurons is a common characteristic of incipient seizure in these two models of acute status epilepticus, and may be a promising element for further study of seizure initiation in models of chronic epilepsy. In addition, our findings highlight the complex involvement of distinct GABAergic interneuron populations in pathological activity in the hippocampal circuit.

## Experimental Procedures

### Animals

All experiments were approved by the Institutional Animal Care and Use Committee of Yale University. Experiments were performed using 2 to 6 month-old PV-Cre (Jackson Laboratory strain #008069) or SOM-Cre (Jackson Laboratory strain #013044) mice crossed with Ai32 (Jackson Laboratory strain #012569) mice for expression of channelrhodopsin-2/EYFP fusion protein in Cre recombinase containing cells (Hippenmeyer et al., 2005; Taniguchi et al., 2011; Madisen et al., 2012). For immunohistological verification of the fidelity of the two Cre lines, they were also crossed to the tdTomato-expressing Ai9 line (Jackson Laboratory strain #007909)(Madisen et al., 2010).

### Acute seizure induction protocol

Recordings were performed in mice anesthetized with .02ml ketamine (80mg/kg)/xylazine (5m/kg) administered by intraperitoneal (IP) injection before they were intubated, mechanically ventilated and then paralyzed using .024mg/g gallamine triethiodide (Sigma). Animals were then fixed in place with a headpost. A 200μm optical fiber was lowered through the cortex until it was ∼200mm above the hippocampal CA1. Recording electrodes were then lowered into CA1 and the optical fiber position was adjusted in small steps. Simultaneous recordings of isolated single units and LFPs were made during spontaneous baseline activity, when ChR2-expressing PV or SOM interneurons were identified with short pulses of light at 473nm, and during periods following chemoconvulsant administration. The preictal period was defined as the time from 350 chemoconvulsant injection until the onset of ictal activity. In all experiments, seizures were induced pharmacologically with IP injection of either 120mg/kg PTZ, a GABA-A receptor antagonist, or 500mg/kg Pilocarpine, a muscarinic AChR agonist.

### Extracellular recordings

All extracellular single-unit, multi-unit, and LFP recordings were made with custom-designed, moveable arrays of tetrodes manufactured in the lab from Formvar-coated tungsten wire (12.5µm diameter; California Fine Wire, Grover Beach CA). Tetrodes were targeted to the CA1 field of the dorsal hippocampus (AP: +1.5-2mm; ML: 1.2-1.75, Franklin and Paxinos, 2001). Signals were digitized and recorded with a DigitalLynx 4SX system (Neuralynx, Bozeman MT). All data were sampled at 40kHz and recordings were referenced to the cerebellum. LFP data were recorded with a bandpass 0.1-9000Hz filter and single-unit data was bandpass filtered between 600-9000Hz.

### Optogenetic manipulations

Light activation of ChR2-expressing cells was performed using a 473nm laser (OptoEngine LLC, Midvale UT). A 200µm fiber was positioned on the cortical surface next to the electrode array and lowered slowly into the cortical tissue directly above dorsal hippocampal CA1. To avoid heating of the brain, we calibrated the light power (<75mW/mm2) during ChR2 unit tracking experiments in order to ensure a mean spike probability of ∼1 spikes per 5 ms light pulse in the targeted population. Real-time output power for each laser was monitored using a photodiode and recorded continuously during the experiment. During baseline periods, we identified ChR2-expressing interneurons using short (5 ms) pulses of blue light, relying on the short latency of ChR2-evoked spikes and the high degree of temporal precision of the evoked spikes. In a subset of experiments (Fig 3-4), we measured the input-output function of ChR2-identified interneurons in response to a calibrated range of light intensities.

### Spike sorting

Spikes were clustered semi-automatically using the following procedure. We first used the KlustaKwik 3.0 software (Kadir, 2013) to identify a maximum of 30 clusters using the waveform energy and energy of the waveform’s first derivative as clustering features. We then used a modified version of the M-Clust environment to manually separate units. Units were accepted if a clear separation of the cell relative to all the other noise clusters was observed, which generally was the case when a conventional metric of cluster separation, isolation distance (ID) (Schmitzer-Torbert et al., 2005) exceeded 20 (Vinck et al., 2015). We further ensured that maximum contamination of the ISI (Inter-spike-interval) histogram did not exceed 0.1% at 1.5 ms. To analyze the firing rates of cells after ictal onset, we manually determined the last point at which cluster separation from the noise was clearly visible. Only cells whose ictal activity could be tracked for at least 60s were included in the ictal period analysis.

### Data analysis tools

All data was analyzed using the Mathworks Fieldtrip toolbox (in particular the Spike toolbox) and custom-made Matlab (The Mathworks, Natick MA) and Igor (WaveMetrics, Lake Oswego, OR) scripts (M. Miri and M. Vinck).

### Analysis of waveform parameters

For each isolated single unit, we computed an average spike waveform for all channels of a tetrode. The waveforms were manually inspected and the channel with the largest peak-to-trough amplitude was used to measure the peak-to-trough duration values (Fig. 1 Supplement 1) as well as mean spike amplitude (Fig. 1 Supplement 5). We also computed the repolarization value of the normalized (between -1 and +1) waveforms at 0.9 ms (similar to Vinck et al., 2015). Non-light-driven units were defined as RS cells by their broad waveform, defined as a repolarization value at 0.9ms smaller than -0.35.

### Seizure detection

Ictal onset was identified by examining hippocampal CA1 LFP recordings, and was defined as the first occurrence of an ictal spike following injection of the chemoconvulsant. This generally corresponded to the LFP trace crossing an absolute z-score value >5 as compared to baseline.Sustained, elevated z-scores were generally observed after ictal onset, and ictal onset was typically coincident with the first ictal spike. We used spectrograms to validate that there were no consistent LFP changes prior to ictal onset (Fig 1A). Spectrograms of LFP power around ictal onset were computed using a wavelet transform with 7 cycles for each frequency and a Hanning taper. LFP power was normalized by dividing by the summed power across the entire trace and taking the base-10 logarithm.

### Definition of analysis periods

Seizure latency varied across mice (Figure S4). For each experiment, the preictal period from injection to ictal onset was therefore divided into four equal periods. To characterize the progressive changes in hippocampal CA1 preictal activity (Figs 1-2), we computed the change in firing activity parameters as compared to baseline for the four preictal periods. For the analysis of spontaneous activity, we only used baseline and preictal periods that did not contain epochs of laser pulses and selected cells with baseline firing rates greater than 0.1Hz. For the analyses of evoked activity in Figures 3-4, all cells were used. For the analysis of RS suppression by ChR2-evoked inhibition, we defined an additional early ictal period as the 60s following ictal onset. For analysis of ictal spiking, we used only the subset of interneurons whose ictal spike activity could be resolved and we divided the 60s ictal period into four periods of 15s each.

### Firing rate and bursting

The mean firing rate per analysis period was computed as the number of spikes in that period divided by the duration of that period in seconds. Changes in the temporal patterning of preictal firing were detected using two metrics. First, we quantified the propensity to engage in irregular burst firing using the coefficient of local variation (LV; Fig 2A), which has been shown to be robust against non-stationarities in firing rates. LV values greater than 1 indicate irregular firing, whereas LV values smaller than 1 indicate sub-Poisson regular firing (Shimokawa and Shinomoto, 2009). Second, we computed the log fraction of ISIs between 2 and 10 ms over the fraction of ISIs between 10 and 100 ms, i.e., Log(ISI_short_/ISI_long_), as in Vinck et al. (2015).

### Spike-field locking and LFP power

Spike-field locking (Fig 2B-C) was computed using the Pairwise Phase Consistency, a measure of phase consistency that is not biased by the firing rate or the number of spikes (Vinck et al., 2012). Spike-LFP phases were computed for each spike and frequency separately by computing the Discrete Fourier Transform of Hanning-tapered LFP segments. These segments 443 had a duration of 7 cycles per frequency. LFP power (Fig. 2 Supplement 4) was computed by dividing the signal into 1s segments and computing the DFT with a Hanning taper. We then averaged the LFP power in the 20-28 Hz band.

### Histology

PV-Cre/Ai9 and SOM-Cre/Ai9 reporter mice were anesthetized with isoflurane and then transcardially perfused through the left ventricle using a peristaltic pump with 0.9% saline solution, followed by 4% paraformaldehyde in 0.1 M phosphate buffer (PB). The brains were removed from the skulls, post-fixed in the same solution at room temperature for 45 mins, and cryoprotected by immersion in 15% and 30% sucrose solutions in PB at 4 °C until they sank.Brains were stored in a −70 °C freezer for further analysis. Coronal sections (20 µm) from hippocampal Ca1 were obtained from each brain using a cryostat (Leica) at −20 °C and collected onto a slide. The sections were carefully washed and treated blocked against non-specific binding with 2% normal goat serum in PB containing 0.1% Triton X-100 (PB-TX) for one hour at room temperature and then incubated with either monoclonal PV (monoclonal; 1:500;Sigma) or SOM (monoclonal; 1:200; Millipore) antibody overnight at 4°C. All sections were again washed in PB-TX and then incubated in an Alexa secondary antibody (1:1000;Invitrogen) for 1 hour at room temperature. Finally, the sections were rinsed in PB and DAPI was added before slides were cover-slipped. Immunostaining and counting was performed on a minimum of three coronal sections from at least three PV-Cre/Ai9 or SOM-Cre/Ai9 animals for each respective condition. Hippocampal analyses were carried out in CA1 and ImageJ (Wayne Rasband, NIH) was used for image processing and counting. To minimize counting bias, we compared sections of equivalent bregma positions (from −1.5 mm to −2.0 mm relative to bregma), defined according to the Mouse Brain atlas (Franklin and Paxinos, 2001).

### Response curves

In a subset of experiments (Fig 3), we measured the input-output function of ChR2-identified 470 interneurons in response to a calibrated range of light intensities. We then made a sigmoid fit of 471 the probability of spiking as a function of light intensity. This sigmoid fit was defined as

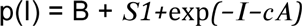

Here, p(I) is the fitted probability of a spike in the 2-15ms following laser pulse onset, I is the laser intensity, S is a scaling factor, c is the c50, and 1/A is the slope. The Rmax was defined as the value of p(I) at the maximum laser intensity tested. We fitted these curves by minimizing the absolute deviation between fit and data (i.e., the L1 norm) using MATLAB’s *fminsearch* function. To avoid finding a local minimum, we randomly chose 64 different starting values for the different parameters and selected the fit that minimized the error across all 64 initializations.

### RS cell inhibition

During a subset of our recordings in PV-Cre/ChR2 and SOM-Cre/ChR2 mice, we simultaneously recorded the activity of local RS cells, defined as described above, and monitored changes in preictal and early ictal inhibition of these units. To quantify the extent to which RS cells were inhibited following the light pulses, we compared their firing rates in the 50ms period post light pulses (FR_post_) with their firing rate in the 200ms prior to light pulses (FR_pre_). We then computed the modulation of RS firing rates (Fig 3C-D) as

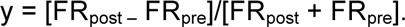

We computed this modulation separately for pulses of medium and high intensity (Fig 4C-D).The medium intensity level was defined as the level at which the simultaneously driven PV cells had, on average, a 50% firing probability. The highest intensity level was the level at which the simultaneously recorded PV cell spiking reached its maximum spike probability.

### Statistical testing

Paired and unpaired Rank Wilcoxon tests were used throughout the manuscript to avoid the assumptions made by parametric statistical tests.

## Acknowledgements

The authors thank M. Higley for comments on the manuscript and H. Blumenfeld for discussion of seizure models. This work was funded by an NSF graduate fellowship and a Ford Foundation graduate fellowship to M.L.M, a Rubicon (NWO) postdoctoral fellowship and a Human Frontiers postdoctoral fellowship to M.V, and a Klingenstein fellowship, a Whitehall grant, an Alfred P. Sloan Fellowship, NIH grant R01 EY022951, and a grant from the Swebilius Foundation to J.A.C.

## Author Contributions

M.L.M. and J.A.C. designed experiments. M.L.M. performed the experiments. M.L.M. and R.P. performed immunohistochemistry. M.L.M. and M.V. analyzed the data. M.M., M.V., and J.A.C. wrote the paper.

